# Evaluation of antiviral treatments for highly pathogenic avian influenza virus infections in feline species

**DOI:** 10.64898/2026.06.09.730954

**Authors:** Alivia C. Ishee, Zhenzhen Zhai, Marek J. Oomens, Ruth N. Collins, Jessica Gomes Noll, Gary R. Whittaker

## Abstract

In 2020, highly pathogenic avian influenza (HPAI) isolates from clade 2.3.4.4b emerged in Europe and spread globally, including in bovine hosts in the USA. Viruses from this clade cause minimal disease in dairy cattle, characterized by decreased milk production but low mortality rates. Infections have also occurred in feline hosts. In contrast to cows, infection of cats (and closely related species, including skunks and foxes) can result in severe neurological signs and mortality. Documented feline H5N1 infections from clade 2.3.4.4.b have a mortality rate of approximately 80% following rapid onset of clinical signs. No antiviral compounds have been tested in an experimental feline model; however, anecdotal clinical evidence suggests early treatment with oseltamivir may improve outcomes in felines with HPAI. Here, we show the *in vitro* efficacy of several influenza inhibitors in feline glial astrocyte (PG-4) and kidney (CRFK) cell culture models using the clade 2.3.4.4.b virus Tx2/24 (H5N1). The neuraminidase inhibitor oseltamivir carboxylate did not effectively inhibit viral replication in either cell line. The cap-dependent endonuclease inhibitor baloxavir exhibited the strongest inhibition of this virus, with EC50 values of 30 nM in PG-4 and 1 μM in CRFK cells. Amantadine and rimantadine, M2 ion channel inhibitors, were unable to completely inhibit viral replication in either cell line at any concentration utilized. The broad-spectrum nucleoside analog GS-441524 demonstrated little to no inhibition of viral replication in either cell line. Additionally, the mutagenic NHC analogs EIDD-1931 and EIDD-2801 successfully inhibited viral replication at the maximum tested concentration of 100 μM but exhibited significant cytotoxicity. Our findings suggest that baloxavir should be considered by veterinary clinicians as the first-line drug of choice when presented with felines or other species infected with HPAI.

## Introduction

Influenza A viruses (IAVs) are orthomyxoviruses known for their capacity to cause highly communicable respiratory disease (1). IAV epidemiology is characterized by both seasonal epidemics driven by human-circulating strains and sporadic pandemic events driven by zoonotic strains (2). Zoonotic IAV strains are thought to originate from wild avian species, including waterfowl and shorebirds (3), and have spread to domestic species such as poultry, swine, and equids, as well as companion animals such as domestic felids and canids (4). IAV genomes comprise 8 segments that code for 11 proteins (5, 6). Concurrent infection of a single cell by two or more IAV strains may result in reassortment, which shuffles the viral genome, leading to genetic diversity and frequent epitope changes (7), facilitating the continued re-emergence of IAV across a diverse range of hosts (8). IAVs are classified by their glycoproteins, hemagglutinin (HA) and neuraminidase (NA), and further classified into clades based on evolutionary relationships (9). Avian influenza strains, including H5N1, are further classified as low-pathogenic avian influenza (LPAI) or highly pathogenic avian influenza (HPAI) based on their HA cleavage site, as the multi-basic cleavage site of HPAI viruses enables cleavage by furin, thereby enabling systemic spread (10). HPAI viruses are of particular interest because they can cause severe disease and mortality in humans through occasional spillovers, although they typically transmit poorly in mammalian species (11).

A new strain of HPAI caused by clade 2.3.4.4 b H5N1 viruses emerged globally in 2020, crossing the Atlantic Ocean into North America in late 2021 (12), and was detected in the United States dairy cow populations in 2024 (13). Clade 2.3.4.4 b H5N1 has infected an unprecedented variety of mammalian species, including minks, foxes, skunks, dolphins, and sea lions (14–17). Particularly troubling are feline infections of clade 2.3.4.4 b viruses, which are characterized by severe neurological signs and high mortality (∼90% for H5N1 clade 2.3.4.4b) (18, 19). As of May 2026, at least 154 documented clade 2.3.4.4b H5N1 infections have been reported in domestic felines in the United States (20). Without comprehensive surveillance, however, many more cases likely remain unreported (21). The primary method of transmission is thought to be contaminated pet food, particularly ‘raw diets’, resulting in exposure to uncooked contaminated poultry (22). Some infected felines are associated with farms with confirmed dairy cattle infections, representing another possible route of exposure (23, 24). While definitive evidence is currently lacking, the history of some feline infections suggests the possible cross-species human-to-cat transmission (23). Importantly, although currently available data indicate it is rare, at least one case of presumptive cat-to-human transmission has been documented (25).

The clinical signs seen in feline H5N1 infections initially include loss of appetite, fever, and lethargy; cats may then rapidly progress to neurological signs, including ataxia, tremors, blindness, and seizures, also accompanied by respiratory signs (dyspnea, tachypnea), and progression to death in most cases (26, 27). There are an estimated 76.3 M owned domestic felines in the United States alone (28), representing a large population of companion animals at risk of infection and capable of transmitting the virus to humans. Such considerations suggest that feline H5N1 infections pose a potential public health threat and must be addressed. Several studies of clade 2.3.4.4 b H5N1 have been conducted *in vitro*, and although ferret and mouse models have been established, antivirals for feline infections have not been directly investigated. A comprehensive panel of available influenza drugs has not been screened for efficacy in a feline model; however, oseltamivir has been used in felines in a single clinical case report (24). In this case report, based on a perceived epidemiological link to HPAI in the area, the attending veterinarian administered oseltamivir to treat two cats presenting with moderate fever and signs of upper respiratory infection (URI). While the cats seroconverted with moderate-to-high H5N1 titers and subsequently recovered, it is not known whether there was direct evidence of a beneficial antiviral effect in these animals. Therefore, in this study, we aim to assess the efficacy of different antivirals used in human and veterinary settings against clade 2.3.4.4.b H5N1 influenza viruses.

We profiled and compared the susceptibility of four feline cell lines and two reference cell lines to infection with Tx2/24. Given the recently elucidated role of astrocytes in facilitating neuroinvasion of clade 2.3.4.4 b viruses in felines (18) and that H5N1 mortality in felines is associated with neurological symptoms, we considered PG-4 (feline glial astrocyte) cells the most relevant cell line to profile for antiviral efficacy and cytotoxicity. We also assessed the antiviral efficacy and cytotoxicity of the selected compounds in CRFK (feline kidney) cell lines to determine if the antivirals are effective in multiple cell lines.

## Materials and methods

### Cell lines

MDCK.1 (CRL-2935), AK-D (CCL-150), CRFK (CCL-94), and PG-4 (S+ L-) (CRL-2032) cells were all obtained from ATCC with the listed catalog numbers. FCWF-CU cells were derived by Dr. Ed Dubovi at the Animal Health Diagnostic Center at Cornell University from FCWF.4 cells from ATCC (CRL-2787). Cal-1 cells were provided by Dr. Diego Diel, Director of Virology at the Animal Health Diagnostic Center at Cornell University. MDCK.1, CRFK, and Cal-1 cells were cultured in Dulbecco’s Modified Eagle’s Medium with 4.5g/L glucose & L-glutamine without sodium pyruvate (Corning, Cat#10-017-CV) with 10% Fetal Bovine Serum, Premium heat-inactivated FBS (Gibco, Cat#A56708-01), and 1% HEPES Buffer 1M (Corning, Cat#25-060-Cl). AK-D cells were cultured in F-12K Nutrient Mixture (Gibco, Cat#21127022) with 10% FBS and 1% HEPES. PG-4 cells were cultured in McCoy’s 5A medium (Gibco, Cat#16600-08-2) with 10% FBS and 1% HEPES. FCWF-CU cells were cultured in Eagle’s Minimum Essential Medium (MEM) with 1.5 g/L sodium bicarbonate, non-essential amino acids, L-glutamine, and sodium pyruvate (Corning, Cat#10-009-CV) with 10% FBS, 10% Corning Nu-Serum™ IV Culture Supplement (Corning, Cat#355504), and 1% HEPES. The H16 antibody was produced by H16-L10-4R5 hybridoma cells obtained from ATCC (HB-65), as described in Tse *et al*. (29). Hybridoma cells were cultured in Hybridoma SFM with L-glutamine (Gibco, Cat#12045-076), beginning at 10% Ultra-Low IgG FBS (Gibco, Cat#16250-078) and weaning to 2.5% FBS over 8 weeks. Cell lines and hybridoma cells were maintained at 37 °C and 5% CO_2_.

### Virus-stocks

Virus stocks were produced by infecting Cal-1 cells with A/dairy cow/Texas/06322424-1/2024 (Tx2/24) (GSAID accession number: EPI_ISL_19155861) provided by Dr. Diego Diel, Director of Virology at the Animal Health Diagnostic Center at Cornell University. Supernatant was harvested and frozen into viral stocks when 50% CPE was observed. Virus stock titers were determined utilizing a TCID_50_ protocol.

### Cell culture

For cytotoxicity and antiviral activity assays, cells were grown to 90% confluency, then split 1:2 using standard media the day prior to plating for infection. Cells were then trypsinized with 0.25% Trypsin, 2.21 mM EDTA 1x without sodium bicarbonate (Corning, Cat#25-052-Cl) and seeded into 96-well white-walled plates (Tissue Culture Plates, white-walled/clear flat bottom, Cell Pro, Cat# TPW0096) at 1 × 10^4^ cells per well. 24 hours post-seeding, plates were used for cell viability or antiviral assays.

### Immunofluorescence assay (IFA) for susceptibility

Cells were plated at a density of 2.5 × 10^4^ cells per well in a 24-well plate (TC Plate, 24-well Standard, Sarstedt, Cat#83.3922) and incubated for 24 hours. The cells were trypsinized and counted to determine the appropriate inoculum concentration to achieve a multiplicity of infection (MOI) of 0.01. Cells were then taken into the BSL-3 facility and infected at an MOI of 0.01 with Tx2/24. Every twelve hours, cells were fixed with 8% paraformaldehyde (Paraformaldehyde solution EM Grade, Electron Microscopy Services, Cat#15714) diluted in PBS (137 mM NaCl, 2.7 mM KCl, 10 mM Na_2_HPO_4_, and 1.8 mM KH_2_PO_4_, pH=7.4) and appropriately decontaminated by submersion. The cells were washed three times with PBS, then permeabilized for 15 minutes at RT with 0.05% Triton X-100 (J.T. Baker, Cat#X198-07) in PBS and blocked for 1 hour at RT in blocking buffer with 1% BSA (Albumin bovine fraction V, Research Products International, Cat#A30075-500.0) in PBS-T (PBS with 0.05% Tween-20 (Fischer Scientific, Cat#BP337-500)). The cells were then stained with H16 anti-Influenza A NP hybridoma supernatant at a 1:10 dilution in blocking buffer overnight. The cells were then washed 3 times with PBS-T and, for visualization, stained with Goat anti-mouse IgG (H+L) Alexa Fluor™ plus 488 (Invitrogen, Cat#A32723) at a 1:1,000 dilution in PBS-T for 1 hour at room temperature, followed by two additional PBS-T washes. DAPI stain was completed by diluting DAPI Fluoromount-G® (SouthernBiotech, Cat#0100-20) 1:5 in PBS and incubating for 15 minutes at RT. The cells were then washed 2 more times with PBS-T and imaged using an ECHO RVL-100M.

### Compounds and reconstitution

Compounds were obtained from MedChemExpress under the following catalog numbers: oseltamivir acid (HY-13318), baloxavir acid (HY-109025A), amantadine (HY-B0402), rimantadine (HY-B0338), GS-441524 (HY-103586), EIDD-1931 (HY-125033), and EIDD-2801 (HY-135853). Upon receipt, lyophilized compounds were centrifuged and reconstituted to a stock concentration of 10 μM for amantadine and baloxavir, and 100 μM for the remaining compounds in freshly opened DMSO (VWR Cat# WN182-10mL). Reconstituted compounds were vortexed briefly, then distributed into single-use aliquots stored at -80 C.

### Cell viability assay

Cytotoxicity was assessed using 96-well plates prepared as previously outlined, incubated at 37 °C in 5% CO2 for 24 hours after treatment. Cell-Titer Glo 2.0 (Promega, Cat #G9242) was then used to assess cell viability according to the manufacturer’s protocol, with a Biotek Synergy H1 plate reader for luminescence detection. Cell viability was then calculated relative to untreated controls and fit to a 3PL curve constrained to a minimum of 0 to estimate CC_10_, the concentration of compound yielding 10% cytotoxicity.

### In-Cell ELISA Antiviral Activity Assay

Cells in 96-well plates were counted via hemocytometer to determine the inoculum concentration. The prepared plates were taken into the BSL-3 facility immediately after treatment and infected at an MOI of 0.1. 24 hours after infection, plates were fixed by submersion in 4% paraformaldehyde in PBS for 30 minutes, then removed from the BSL-3 facility.

Cells were permeabilized with 0.5% TritonX-100 in PBS for 15 minutes at RT and blocked with 2% Non-fat Dry milk (Lab Scientific bioKEMIX, Cat#M0841) (w/v) in PBS-T for 1 h at room temperature (RT). Cells were stained with the mouse monoclonal pan-influenza A NP antibody H16, derived from hybridoma supernatant, at a 1:10 dilution overnight in blocking buffer, then washed 3x with PBS-T. Secondary antibody staining was performed with goat anti-mouse IgG HRP (Invitrogen, Cat#31430) reconstituted to 0.8 mg/mL at a 1:10,000 dilution in PBS-T, followed by 4x PBS-T washes. Cells were treated with 1-Step™ Ultra TMB-ELISA (Thermo Scientific, Cat#34029) for 15 minutes at RT, then quenched with 0.5 M H2SO4 (Thermo Scientific, Cat#12424-0010). Plates were then read with a Biotek Synergy H1 plate reader for absorbance at 450 nm and 620 nm.

Percent infection was calculated as a function of 0% infection (uninfected, untreated control) and 100% infection (infected, untreated control) and fit to a logarithmic response curve to interpolate EC_50_ (50% viral replication). Calculations and graphing were carried out in GraphPad Prism 11. The selectivity index (SI) was calculated for each tested drug with an EC50 within the tested range for each cell line. Since CC_50_ values were not observed in the tested range, CC_50_ for SI calculation is assumed to be the maximum tested concentration, and SI is reported as a number > than the maximum concentration/EC_50_.

## Results

### Susceptibility of feline cell lines to H5N1 infection

Cell line susceptibility was assessed by infecting confluent monolayers of Cal-1 (bovine uterine) and MDCK.1 (canine kidney) cells for comparison. Feline cell lines, including AK-D (feline epithelial lung), CRFK (feline kidney), FCWF-CU (feline macrophage-like), and PG-4 (feline glial astrocyte) cells, were also profiled for susceptibility. All susceptibility testing was completed at an MOI of 0.01, and viral replication in cells was imaged using IFA every 12 hours (Figure 1). All tested cell lines supported infection with Tx2/24, with Alexa Fluor 488 signal increasing over time, corresponding to increased detection of influenza A nucleoprotein. The increased signal indicated that all tested cell lines are both susceptible and permissive to infection with Tx2/24.

**Figure 1).**
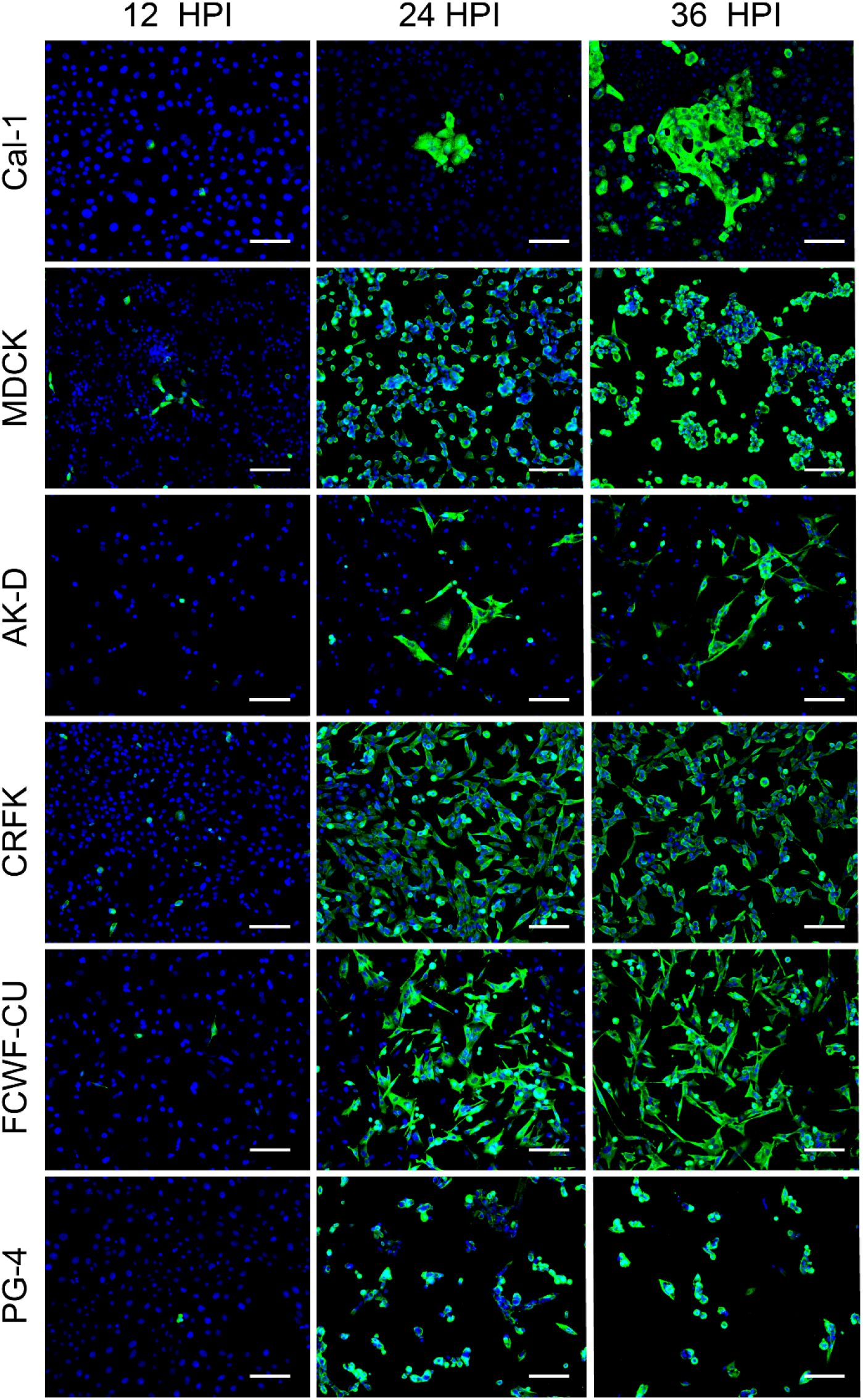
Susceptibility of feline and reference cell lines to infection with H5N1. Cal-1 (bovine uterine), MDCK (canine kidney), AK-D (feline epithelial lung), CRFK (feline kidney), FCWF-CU (feline macrophage-like), and PG-4 (feline glial astrocyte) cell lines infected with Tx2/24 at 12, 24, and 36 hpi. Viral replication was visualized using H16 (pan-influenza A NP mAb) and Alexa Fluor 488 Plus, with blue DAPI staining. All scale bars are 130 µm.

### Antiviral efficacy and cytotoxicity of compounds in PG-4 cells

Compounds were assessed for antiviral activity and cytotoxicity using log-fold dilutions from stock solutions at 100 mM for oseltamivir-carboxylate, rimantadine, EIDD-1931, EIDD-2801, and GS-441524, and at 10 mM for baloxavir-acid and amantadine. Maximum stock concentrations were determined by compound solubility in DMSO, and all initial dilutions were 1:1000, yielding a maximum DMSO concentration of 0.01%. Antiviral testing was performed at an MOI of 0.1 to recapitulate active infection and assess treatment rather than prophylaxis.

In PG-4 cells, at maximum concentration, all tested compounds except GS-441524 demonstrated a statistically significant reduction of viral replication compared to the maximum DMSO concentration as assessed by Welch’s t-test with p < 0.05 (Figure 2A). Only baloxavir and EIDD-1931 demonstrated statistically significant reduction in cell viability at maximum concentration in comparison to DMSO, as assessed by Welch’s t-test with p < 0.05 (Figure 2B).

**Figure 2).**
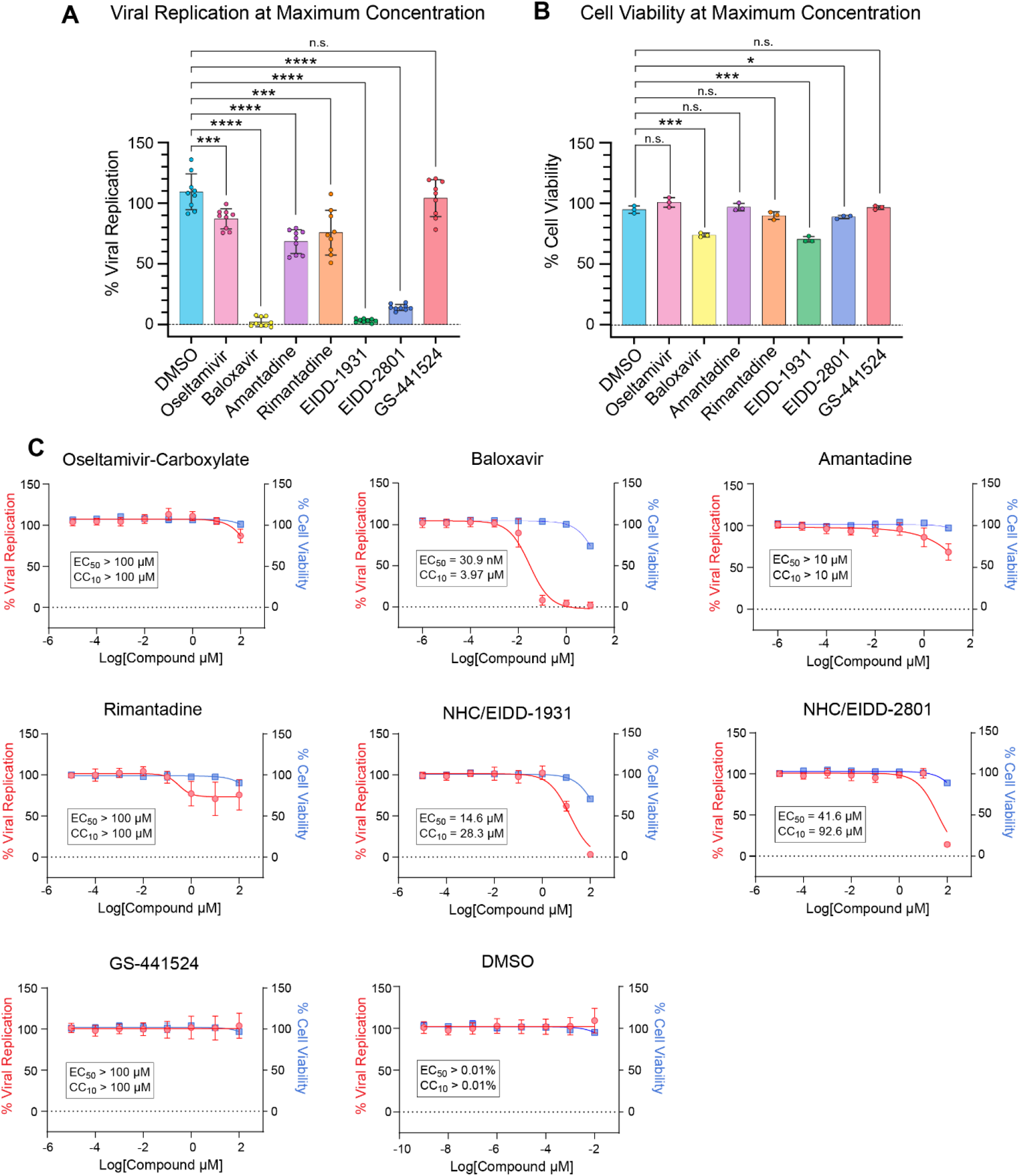
Antiviral activity and cytotoxicity of antivirals in PG-4 cells infected with H5N1. A) Viral replication at maximum concentrations of tested drugs (0.01% DMSO; 100 µM oseltamivir-carboxylate, amantadine, rimantadine, EIDD-1931, EIDD-2801, and GS-441524; 10 µM baloxavir, and amantadine, 100 µM) as determined through in-cell ELISA 24 hpi with MOI 0.1 using Tx2/24. B) Cell viability at maximum concentrations of tested drugs assessed by Cell-Titer-Glo 2.0 24 hours after treatment C) Dose response of antiviral activity and cytotoxicity of compounds (n.s.P > 0.05, *P ≤ 0.05, **P ≤ 0.01, ***P ≤ 0.001, ****P ≤ 0.0001)

### Oseltamivir-carboxylate does not inhibit H5N1 replication in PG-4 cells

The current first-line drug for avian influenza infections in humans is oseltamivir (Tamiflu®) (30). Oseltamivir is a neuraminidase inhibitor that functions by preventing the release of influenza A virions by preventing their cleavage from sialic acid, and it is FDA-approved for the treatment of uncomplicated influenza (31, 32). The recommended oral course of treatment is twice daily for 5 days (33). Oseltamivir (oseltamivir-phosphate) is an oral prodrug metabolized in the liver by carboxylesterase 1 (CES1) prior to circulation in the bloodstream as the active drug, oseltamivir-carboxylate (34). In the absence of a system to metabolize the prodrug, we tested the active drug, oseltamivir-carboxylate in cell culture. Oseltamivir carboxylate demonstrated no statistically significant reduction in H5N1 infection in PG-4 cells at maximum concentration, and there was no dose-dependent response at subsequent dilutions (Figure 2C). EC_50_ and CC_10_, therefore, lie outside the tested concentrations and cannot be reported.

### Baloxavir strongly inhibits H5N1 replication in PG-4 cells

Baloxavir-marboxil (Xofluza®), the most recent FDA-approved influenza-specific antiviral, targets the cap-dependent endonuclease activity of the viral polymerase, PA (35, 36). The recommended course of treatment for humans is a single oral dose (35). Baloxavir-marboxil is an orally available prodrug that is rapidly metabolized by uridine monophosphate glucuronosyl transferase 1A3 (UGT1A3) (35). In the absence of a system to metabolize the prodrug, we tested the active form of the drug, baloxavir-acid in cell culture. Baloxavir acid significantly reduced viral replication and cell viability at the maximum concentration. Baloxavir strongly inhibited H5N1 replication in PG-4 cells in a dose-dependent response (Figure 2C), achieving 100% inhibition at higher concentrations and an EC_50_ value of 30.9 nM and a CC_10_ value of 3.97 µM (Table 1). The selectivity index (SI) was calculated as >3,236 by (maximum tested concentration)/EC50, since CC_50_ was not observed.

**Table 1).** EC_50_, CC_10_, and SI of tested compounds in PG-4 and CRFK cell lines.

| Compound | EC<sub>50</sub> (μM) |  | CC<sub>10</sub> (μM) |  | SI |  |
| --- | --- | --- | --- | --- | --- | --- |
|  | PG-4 | CRFK | PG-4 | CRFK | PG-4 | CRFK |
| **Oseltamivir** | >100 | >100 | >100 | >100 | NA | NA |
| **Baloxavir** | 0.0309 | 0.0320 | 3.97 | 3.13 | >3,236 | >3,125 |
| **Amantadine** | >10 | 1.14 | >10 | 3.33 | NA | >87.7 |
| **Rimantadine** | >100 | 0.133 | >100 | 99.0 | NA | >751 |
| **EIDD-1931** | 14.6 | 11.5 | 28.3 | 28.3 | >6.84 | >8.70 |
| **EIDD-2801** | 41.6 | 55.0 | 92.6 | 68.5 | >2.40 | >1.81 |
| **GS-441524** | >100 | >100 | >100 | 42.5 | NA | NA |
| **DMSO** | >0.01% | >0.01% | >0.01% | >0.01% | NA | NA |

### Amantadine and rimantadine weakly inhibit H5N1 replication in PG-4 cells

The amantadates, amantadine (Symmetrel®), and its derivative rimantadine (Flumadine®), are first-generation influenza antivirals that specifically inhibit the M2 ion channel of influenza A viruses, preventing proper genome uncoating and viral egress in avian influenza strains (37, 38). Neither amantadine nor rimantadine is currently recommended by the CDC for the treatment of seasonal influenza due to the rapid emergence of resistant strains (39). Despite widespread resistance to seasonal influenza and occasional resistance to avian influenza strains, H5N1 strains from clade 2.3.4.4 b remain broadly susceptible to treatment with amantadates (40, 41). Amantadine and rimantadine demonstrated statistically significant reductions in viral replication, but neither completely inhibited replication at maximum concentrations of 10 µM and 100 µM, respectively. Neither compound exhibited statistically significant reductions in cell viability at these concentrations. Neither compound reached 50% inhibition of viral replication or 10% reduction in cell viability, so an EC_50_ or CC_10_ value could not be calculated (Figure 2C).

### GS-441524 does not inhibit H5N1 replication in PG-4 cells

GS-441524 is an adenosine nucleoside analog prodrug that is metabolized to the same bioactive molecule as remdesivir, which targets the viral RNA-dependent RNA polymerase (RdRp) required for viral genome replication, with delayed chain-termination activity (42). Due to the highly conserved nature of viral RdRps, many nucleoside analogs have broad-spectrum activity against a range of RNA virus families, including remdesivir (Veklury®), which exhibits antiviral activity against a range of viruses, including coronaviruses, paramyxoviruses, orthomyxoviruses, and filoviruses (42–45). While remdesivir has shown limited impact for treatment in human patients, GS-441524 has demonstrated great success in treating feline infectious peritonitis (FIP) caused by feline coronavirus and received an enforcement exemption from the FDA in 2024, enabling caretakers to access the medication through compounding pharmacies (46, 47). Given the popularity and widespread use of the drug among veterinary clinicians, and early data supporting its activity against orthomyxoviruses (42), we tested it against H5N1. However, GS-441524 did not demonstrate a statistically significant reduction in viral replication at the highest tested concentration (100 µM) (Figure 2A).

### NHC nucleoside analogs inhibit H5N1 replication in PG-4 cells

Similar to GS-441524, N^4^-hydroxycytidine (NHC) analogs, EIDD-1931 and EIDD-2801 (molnupiravir (Lagevrio®)) have recently been repurposed in veterinary medicine for the treatment of FIP and other viruses (48). It is worth noting that neither NHC analog possesses FDA approval for influenza treatment nor exemption from federal enforcement, as the influenza-specific antivirals or GS-441524 do. Further, NHC analogs inhibit viral replication by inducing lethal mutagenesis beyond the virus’s tolerated threshold, resulting in nonfunctional mutants, which differs from the mechanism of action of GS-441524 (49). EIDD-1931 and EIDD-2801 exhibited statistically significant reduction of H5N1 replication in PG-4 cells, with EIDD-1931 achieving 100% inhibition at the highest concentration tested (Figure 2A). EIDD-1931 exhibited a statistically significant reduction in cell viability at the maximum tested concentration, while EIDD-2801 did not (Figure 2B). Both compounds reached 50% inhibition of viral replication and 10% loss of cell viability in the tested range (Figure 2C). Utilizing a 3PL curve constrained with a bottom value of 0, the EC_50_ was calculated to be 14.6 µM and 41.6 µM, and the CC_10_ was 28.3 µM and 92.6 µM for EIDD-1931 and EIDD-2801, respectively, with respective selectivity indices of >6.84 and >2.40 (Table 1).

### Feline kidney cell line testing recapitulates findings in feline glial astrocytes

Many of the tested compounds are prodrugs that require host liver metabolism to be converted into their active forms. When possible, we administered the active forms of the drugs (oseltamivir carboxylate and baloxavir acid). The activation of nucleoside analogs depends on intracellular metabolism in the target tissue (50), and the expression profiles of the metabolic machinery vary across tissue types. Therefore, we also tested all compounds in CRFKs, a feline kidney cell line (Figure 3). The results were consistent with those in PG-4 cells and are summarized in Table 1.

**Figure 3).**
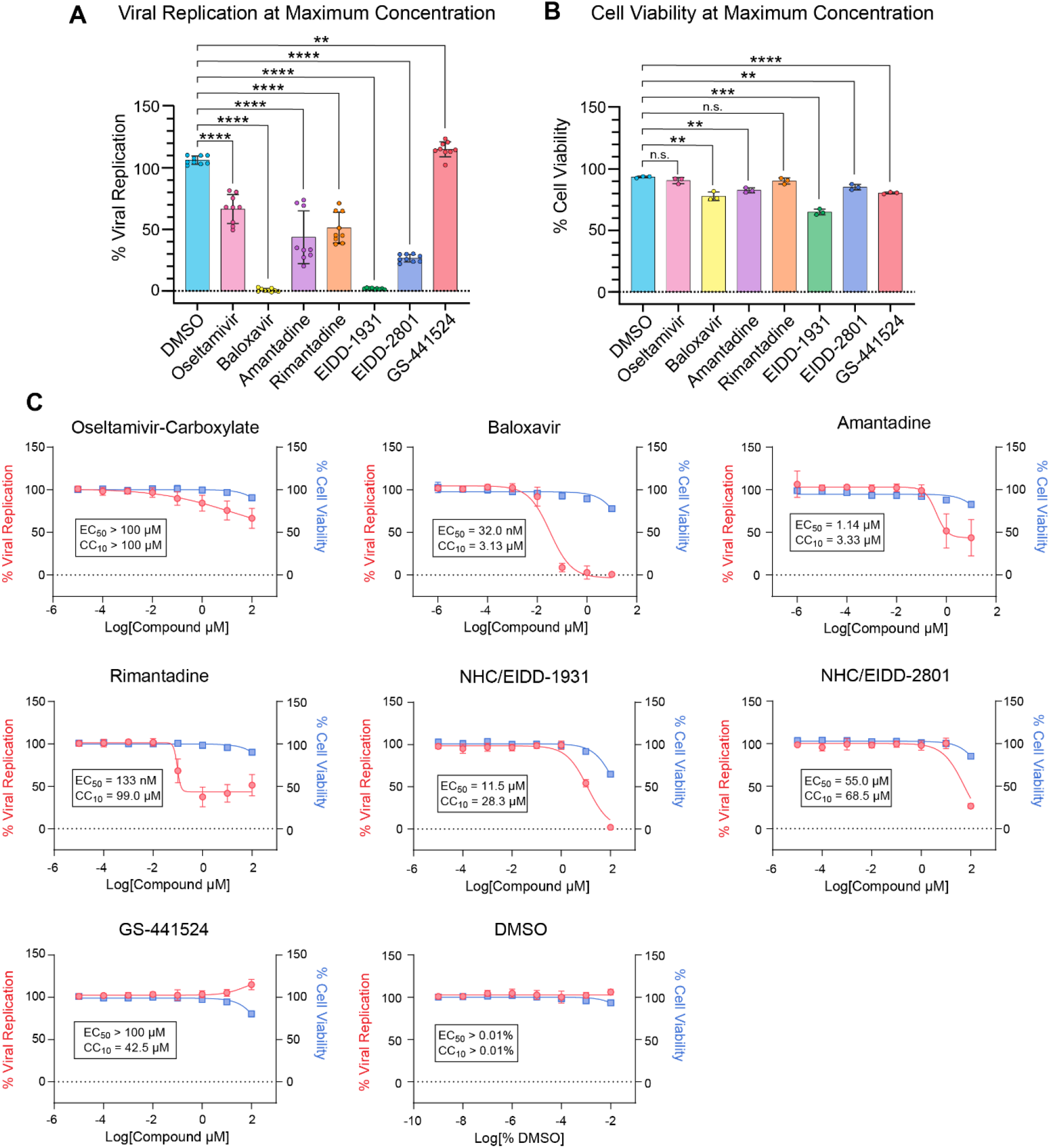
Antiviral activity and cytotoxicity of antivirals in CRFK cells infected with H5N1. A) Viral replication at maximum concentrations of tested drugs (0.01% DMSO; 100 µM oseltamivir-carboxylate, amantadine, rimantadine, EIDD-1931, EIDD-2801, and GS-441524; 10 µM baloxavir, and amantadine, 100 µM) as determined through in-cell ELISA 24 hpi with MOI 0.1 using Tx2/24. B) Cell viability at maximum concentrations of tested drugs assessed by Cell-Titer-Glo 2.0 24 hours after treatment C) Dose response of antiviral activity and cytotoxicity of compounds (n.s.P > 0.05, *P ≤ 0.05, **P ≤ 0.01, ***P ≤ 0.001, ****P ≤ 0.0001)

## Discussion

In early 2021, clade 2.3.4.4.b H5N1 viruses were detected in North America, continuing their global spread across six continents (51–53). This pervasive spread has led to the establishment of new mammalian reservoirs, notably cattle and felines (13, 54). Domestic feline clade 2.3.4.4.b H5N1 infection is associated with severe neurological signs and a high mortality rate (55– 57). While severe neurotropic influenza in a mammalian species is, in and of itself, concerning, the fact that domestic felines can act as a vector linking humans to reservoir species such as cattle and birds is of particular public health concern. This risk is underscored by serological evidence of a presumed cat-to-human H5N1 transmission event involving a veterinarian and an infected domestic feline (25). While several antiviral treatments are FDA-approved for use in humans with H5N1, there is currently no therapeutic guidance for infected felines.

Our data demonstrate that baloxavir acid was the most effective monotherapy for reducing H5N1 replication in feline cell lines, achieving an EC_50_ of 30.9 nM in PG-4 cells. This finding is consistent with current H5N1 clade 2.3.4.4.b *in vivo* models, in which baloxavir treatment reduced mortality in both mice and ferrets, which also exhibit neurological signs upon infection with H5N1 (40, 58–60). In these studies, survival outcomes were generally improved by increasing the dose of baloxavir from 0.1 mg/kg to 10 mg/kg, as demonstrated by Andreev *et al*., and by increasing the number of doses, as shown by Jones *et al*. (40, 59). Interestingly, the worst *in vivo* survival outcomes occurred in studies by Liu *et al*., in which baloxavir was orally administered as the prodrug, baloxavir marboxil, compared with other studies in which the active form of the drug, baloxavir acid, was injected subcutaneously. Previous research suggests that while ferrets poorly metabolize baloxavir marboxil into baloxavir acid, BALB/c mice are readily able to metabolize baloxavir marboxil into baloxavir acid (58, 61, 62).

The pharmacokinetics, pharmacodynamics, and safety of baloxavir marboxil in felines are currently unknown and warrant investigation. The results of such studies would provide important information on potential adverse drug reactions and the best formulation to administer to H5N1-infected felines, depending on their metabolic capacity. Until such studies can be conducted, subcutaneous baloxavir acid treatment may be more effective than oral baloxavir marboxil because it bypasses in-host metabolism.

Beyond reducing mortality in felines, treatment with baloxavir may also help reduce the potential for cross-species transmission of H5N1 between felines and humans. Treatment of H5N1-infected humans with baloxavir led to fewer incidences of contact transmission when compared with a placebo treatment, likely due to a rapid decrease in viral shedding following initiation of treatment (63, 64). A decrease in H5N1 viral shedding after baloxavir treatment was also observed in a ferret model, in all baloxavir acid dosing regimens at the 3, 5, and 7 days post-infection time points, further corroborating that baloxavir treatment may reduce shedding in H5N1-infected felines, thereby decreasing the likelihood of feline-to-human transmission (59).

Although feline H5N1 infections appear to progress rapidly once clinical signs are observed, the infection kinetics and optimal treatment windows remain unknown. In an H5N1 ferret model described in Kiso *et al*., a 48-hour delay in baloxavir marboxil therapy resulted in 100% mortality (55, 56, 65). While 100% of ferrets treated 1 hpi survived in this study, only 40% of ferrets treated 24 hpi survived, with most ferrets succumbing to infection on 13 dpi, 7 days after cessation of treatment. These findings suggest that a longer course of treatment in felines with delayed initiation may improve survival outcomes, as observed in the ferret model by Jones *et al*. (59, 65). Further, baloxavir may be most effective as a prophylactic in preventing infection and mortality in cats with a known exposure to clade 2.3.4.4.b H5N1 viruses; however, treatment should be given judiciously out of concern for resistance mutations, which are known to occur in both human populations and single-dose animal models (59, 66).

Neuraminidase inhibitors (NAIs), such as oseltamivir, are first-line treatment options for influenza infections (both seasonal and avian) in humans, supported by established practice and robust safety data. In addition to oseltamivir, zanamivir (Relenza®) is another neuraminidase inhibitor, not included here because it is not widely used, in part due to its oral-nasal route of administration. In one case series, oseltamivir was presumed to have reduced mortality associated with H5N1 in domestic felines. In a single household, two untreated symptomatic felines died, while the two subsequently treated felines that were given 15 mg of oseltamivir twice daily for ten days survived and were later found to be seropositive for H5N1 (24). Importantly, the lack of a negative control group and controls for other confounding factors in this report limited the ability to establish a causal relationship between oseltamivir treatment and outcomes.

Although presumed to be high, the mortality rate of H5N1 in naturally infected felines remains unknown due to limited surveillance for subclinical cases, likely inflating the perceived mortality rate (21). In our study, we did not find a statistically significant reduction in viral replication even at the highest tested oseltamivir concentration, indicating that it likely had little effect in our system. Recent ferret and mouse models corroborate the notion that oseltamivir treatment did not result in a statistically significant reduction in weight loss or viral shedding compared with vehicle-only controls (40, 59, 60). In humans, oseltamivir tends to be more effective when administered early in infection, preventing viral replication when viral titers are low, and is less effective once infection is well established, when viral replication peaks (67). In the study by Gomez *et al*., treatment was initiated when clinical signs were mild, which may explain why oseltamivir was effective in preventing severe symptoms and mortality in treated domestic felines but remained ineffective in experimental ferret and mouse models. Additionally, some mouse and ferret models suggest that a combination therapy of baloxavir and oseltamivir may yield better clinical outcomes, reduce viral shedding, and may similarly be effective in felines (58, 60).

M2 ion channel inhibitors amantadine and rimantadine exhibited partial inhibition of viral replication in our study, but at the concentrations tested, 100% inhibition was not achieved. This finding agrees with previous ferret models in which amantadine did not result in an improvement of clinical outcomes of ferrets infected with H5N1 (40). Historically, influenza viruses have also readily developed resistance to M2 ion channel inhibitors after treatment, as demonstrated by high levels of amantadine resistance among H5N1 avian viruses isolated from Northern China following extensive amantadine use in the early 2000s (68). For this reason, M2 ion channel inhibitors are not recommended for widespread treatment of influenza and would likely be poor drug candidates in infected domestic felines (41). The broad-spectrum nucleoside analog GS-441524, widely used to treat FIP, was tested because of its conserved mechanism of action and established safety and efficacy in feline populations. It is worth noting that we chose not to test favipiravir, another nucleoside analog expressly approved for the treatment of influenza in Japan (69), due to its lack of availability in the United States. GS-441524 at the highest tested concentration of 100 µM yielded no statistically significant reduction in viral replication in feline glial astrocytes, indicating that it does not inhibit Tx2/24 and should not be considered for treatment in felines with H5N1.

NHC nucleoside prodrugs, including EIDD-1931 and EIDD-2801 (molnupiravir), have been evaluated for efficacy against human, avian, and swine-origin influenza A virus strains (70). Both EIDD-1931 and EIDD-2801 are nucleoside prodrugs that metabolize into the same active antiviral compound, EIDD-2061; both of these have been widely used in veterinary medicine as alternative treatments for FIP in felines (49, 71). Given their popularity and efficacy as influenza treatments, we wanted to assess their efficacy against Tx2/24 in PG-4 cells. Both compounds inhibited viral replication at the highest tested concentration, 100 µM. However, complete inhibitory activity was observed only at the highest concentration, which is unlikely to be reached in the serum of a living feline. Further, the ‘therapeutic window,’ defined as the concentrations that completely inhibited viral replication without cytotoxicity, was nonexistent. The mechanism of action of NHC nucleoside analogs is lethal mutagenesis (49). This treatment regimen can also generate new viral mutants. It may contribute to host-cell mutagenesis over prolonged treatment (72, 73). Although this claim has been disputed, such considerations raise further concerns about the safety of this drug in felines and caution against its use from a One Health perspective (74).

In conclusion, we demonstrate that baloxavir acid is highly effective in reducing viral replication of Tx2/24 in feline cell lines. Pending demonstration of safety and pharmacokinetics/pharmacodynamics in cats, we feel that consideration of this drug as the first-line treatment for symptomatic H5N1-infected felines is a reasonable choice, especially considering the poor and grave prognoses of felines in this cohort. Our work is extensively corroborated by findings from ferret and mouse models, in which baloxavir consistently outperforms all other FDA-approved influenza antivirals. Baloxavir has also been shown to reduce viral shedding and transmission in animal and human models, potentially helping prevent further feline-to-human transmission.

## Acknowledgements

We thank the Diel Lab for providing us with Tx2/24 H5N1 stocks and Cal-1 cells for viral propagation.

All work with H5N1 HPAI was completed at the BSL-3 facility housed in the Animal Health and Diagnostic Center at Cornell University. We thank the BSL-3 staff, Paul Jennette, and Nicole Kushner for their training and support.

Funding was provided by the Cornell Feline Health Center (CFHC) Feline Viral Surveillance Consortium. DOI: 10.2460/ajvr.26.05.0208

Thank you to Dr. Bruce Kornreich and Sue Holster for reviewing this manuscript for clarity and accuracy in relation to veterinary sciences. Special thanks to the Whittaker Lab for their support and guidance in conducting experiments, analyzing data, and providing general support.

